# Apoplast metabolomics reveals that *plant-pathogen* crosstalk is modulated by nitrogen supply

**DOI:** 10.1101/2024.11.11.622918

**Authors:** Roua Jeridi, Antoine Davière, Sylvie Jolivet, Marco Zarattini, Gilles Clement, Marie-Christine Soulie, Ahmed Landoulsi, Mathilde Fagard

**Affiliations:** Université Paris-Saclay, INRAE, AgroParisTech, Institute Jean-Pierre Bourgin for Plant Sciences (IJPB), 78000, Versailles, France; Laboratoire des risques liés aux stress environnementaux, lutte et prévention, UR17ES20, Faculté des Sciences de Bizerte, Université de Carthage, Tunisia; Université libre de Bruxelles (ULB), Faculté de Science, Interfaculty School of Bioengineers, Brussels, Belgium; Sorbonne Université, UPMC Université Paris 06, UFR 927, Paris 75005, France

**Keywords:** apoplast, nitrogen limitation, multistress, virulence, *hrp*, metabolome, *Arabidopsis thaliana*, *Erwinia amylovora*

## Abstract

In the present study, we analyzed the role played by the apoplast in the crosstalk between biotic and abiotic stress conditions. In particular, we studied the crosstalk between nitrogen (N) limitation and infection of the model plant *Arabidopsis thaliana* by *E. amylovora*, an apoplastic bacterium. Our previous findings indicated that low N (LN) conditions increase *E. amylovora in planta* titers and expression of virulence factors. In this work, we extracted the apoplast wash fluids (AWF) from plants grown under low N or high N (HN) conditions and applied them to bacteria *in vitro*. We observed that LN-AWF induced stronger virulence gene expression than HN-AWF. Metabolomic analysis of both apoplast extracts revealed the presence of common metabolites, however, their proportions were distinct, indicating a direct effect of N availability on apoplast content. Interestingly, changes in the apoplast metabolite proportions were also observed early after bacterial infection, but only in plants grown under LN conditions. To evaluate the effect of single metabolites on virulence gene expression, we selected 43 metabolites and observed that 29 of them were activators whereas two, GABA and citrate, acted as repressors. This study shows that environmental constraints, such as N availability, impact plant-pathogen interactions by altering the apoplastic content.

## INTRODUCTION

In plants, the apoplast compartment is the space outside the plasma membrane that includes the cell wall, a network of interconnected extracellular spaces, and the fluid that moves within the walls and the plasmalemma. This fluid is known as apoplast fluid which accounts for around 4-11% of the total leaf fresh weight (Clarkson 2007). The apoplast is involved in many different physiological responses, including the transport of water and solutes throughout the plant, thus playing a key role in the life of plants. It contains large numbers of molecules such as cell wall constituents, proteins, and both primary and specialized metabolites (Clarkson 2007). Although poorly characterized, the apoplastic fluid is known to contain small metabolites, inorganic ions and proteins, the concentration of which depends on the plant species, circadian rhythm, plant age and growth conditions (Green *et al*. 2020; Lohaus *et al*. 2001; Rico *et al*. 2011; Solomon & Oliver 2002; Tejera *et al*. 2006). Furthermore, the apoplast is thought to play a key role in the adaptation of plants to their environment since its content is modulated by biotic and abiotic stresses such as high or low temperatures, water deficiency or submergence, darkness or high light, nutritional deficiency and mechanical stress (Hoson, 1998; López-Milan *et al*. 2001).

Bacterial pathogens that enter the leaf tissue through hydathodes, wounds and stomata are rapidly in contact with the apoplast which constitutes their ecological niche providing a source of nutrition for these pathogens (Rico & Preston 2010). This compartment is the place where early plant-pathogen interactions take place and where the pathogen faces the first line of defense (proteins and metabolites) secreted by plant cells. It is also in this compartment that bacteria perceive initial plant-derived signals to trigger their pathogenesis program (Rico & Preston 2010). The type 3 secretion system (T3SS) is a widespread virulence factor that allows bacteria to inject protein effectors in the plant cytoplasm to modify a range of host targets and attenuate defense (Schreiber, Chau-Ly & Lewis 2021). The T3SS, encoded by hypersensitive response and pathogenicity (*hrp*) genes, is one of the major virulence determinants of gram-negative plant pathogenic bacteria such as *Pseudomonas syringae*, *Ralstonia solanacearum, Xanthomonas* species, and *Erwinia amylovora.* The expression of *hrp* genes is induced *in planta* but the molecular factors that determine their expression are not well known (Asif, Xie & Zhao 2024). *In vitro* studies have shown that *hrp* expression is influenced by environmental conditions such as temperature (Huot *et al*. 2017), pH (Brencic & Winans 2005) and primary metabolite concentration (Stauber, Loginicheva & Schechter 2012;). Furthermore, *hrp* gene expression is induced by biochemical and physiological signals that are characteristic of the leaf apoplast, such as low pH and low nitrogen (N) (Huynh, Dahlbeck & Staskawicz 1989; Wei, Sneath & Beer 1992). In addition, bacterial *hrp* genes are generally more highly expressed in minimal medium, such as M9, compared to rich medium such as Luria Broth (LB) or King B (KB) (Wei *et al*. 1992; Rahme *et al*. 1992; Preston 2000; Rico & Preston 2008; Wang *et al*. 2020). However, the effect of apoplast content fluctuations on bacterial *hrp* gene expression *in plant*a is currently not known.

Our previous work carried out on the *E. amylovora* - *A. thaliana* interaction showed that both bacterial growth and necrotic lesions on leaves require a functional T3SS (Degrave *et al*. 2008). In the context of this non-host interaction, the expression of *E. amylovora* virulence genes is limited and hinders its multiplication in the first hours of infection (Farjad *et al*. 2021). Interestingly, we showed that low N supply favors early *in planta* expression of *hrp* genes and multiplication of *E. amylovora* (Farjad *et al*. 2021). Since *E. amylovora* is localized in the apoplast during the early stages of *A. thaliana* leaf infection (Degrave *et al*. 2008), we hypothesized that differences in *hrp* gene expression could be due to variations in apoplastic metabolite content under different N supply conditions.

In this paper, we extracted apoplast wash fluid (AWF) from plants grown under low N (LN) and high N (HN) and tested their effect on the induction of bacterial *hrp* genes *in vitro*. Interestingly, the activation of virulence genes was observed only following treatment with LN-AWF. Gas chromatography-mass spectrometry (GC-MS) analysis revealed the presence of common metabolites in both apoplast extracts, however, different concentration levels were observed. Selected metabolites were chosen and their *hrp-*inducing or -repressing capacities were studied. Interestingly, we identified two metabolites, GABA and citrate, with repressor capacity. Altogether, our data show that the abiotic environment impacts plant metabolic content both before and after bacterial infection, thus affecting the outcome of biotic interactions.

## RESULTS

### Apoplast wash fluid (AWF) extraction from plants grown in LN and HN

To determine the effect of nitrogen (N) supply on apoplast content, we followed the vacuum infiltration-centrifugation method, which allows the extraction of metabolites compatible with mass spectrometry analysis (Lohaus *et al*. 2001). Col-0 *A. thaliana* plants were grown respectively under 2 mM NO_3_ ^-^ (LN) and 10 mM NO3^-^ (HN) and AWF was extracted after five weeks. A minimum of three independent experiments were performed. We infiltrated leaves with Milli-Q water at relatively low vacuum pressure (25 KPa), and used an intermediate centrifugation force (1000 g). Since the process of infiltration dilutes the apoplastic fluid, it was necessary to determine the extent of this dilution. The dilution factor, calculated by measuring indigo carmine dye dilution, was 1.06 ± 0.10 for LN-AWF and 2.08 ± 0.10 for HN-AWF. To detect cytoplasmic contamination in our apoplast extracts, we measured cytoplasmic markers such as the activity of the enzyme malate dehydrogenase (MDH) and chlorophyll content either in AWF or leaf extracts from plants growth under low or high N supply. MDH levels were estimated at 0.04 ± 0.0062 U/ml in LN-AWF and 0.02 ± 0.0025 U/ml HN-AWF, concentration significantly lower as compared to MDH activity measured in leaf extracts which were respectively 212.54 ± 7.8 and 328.22 ± 7.4 U/ml in plants grown in LN and HN. These data are consistent with previously published reports and show that only 0.02 % and 0.01 % of our LN- and HN-AWF extracts, respectively, correspond to cytoplasmic contamination (Floerl *et al*. 2012; O’Leary *et al*. 2016). These results confirm that the vacuum infiltration method used in this study is reliable, reproducible, and generates minimal cytoplasmic contamination.

### LN-AWF induces *hrp* gene expression and increases the fitness of *E. amylovora*

To test the hypothesis that N availability affects *E. amylovora hrp* expression through modifications of the apoplast content, we incubated *E. amylovora* in LN-AWF and HN-AWF and analyzed the ability of both apoplast extracts to induce *hrp* expression *in vitro*. We used the *E. amylovora* 6023 strain which contains the *lacZ* reporter gene, encoding the ß-galactosidase enzyme, upstream of *the hrp* cluster (Barny *et al*. 1990). Expression of *lacZ* was monitored by measuring ß-galactosidase activity, quantified using an artificial substrate, o-nitrophenyl-ß-D-galactopyranoside (ONPG) (Griffith 2002). As a control, the reporter strain was cultured either in the inducing medium (IM), a minimal culture medium composed of M9 salts, nicotinic acid and galactose, which is known to induce *hrp* expression (Gaudriault, Malandrin, Paulin & Barny 1997), or in Luria-Bertani (LB), a culture medium unable to induce *hrp* expression. In addition, we also monitored *hrp* expression of bacteria incubated in whole leaf extracts of plants grown under low N (LN-L) or high N (HN-L) conditions (Figure 1). As expected, incubation in IM led to significant activation of *hrp* gene expression as compared to LB, a rich medium. When bacteria were incubated with LN-L, *hrp* gene expression was similar to that observed in IM-treated bacteria (Figure 1). On the other hand, slightly lower *hrp* gene expression was observed following HN-L treatment. Although both AWF extracts were able to significantly induce *hrp* expression in *E. amylovora*, LN-AWF was the strongest inducer among all the tested extracts with a 4-fold increase compared to HN-AWF (Figure 1). These results suggest that low nitrogen conditions make the plant apoplast a more favorable environment for *hrp* gene expression.

**Figure 1:**
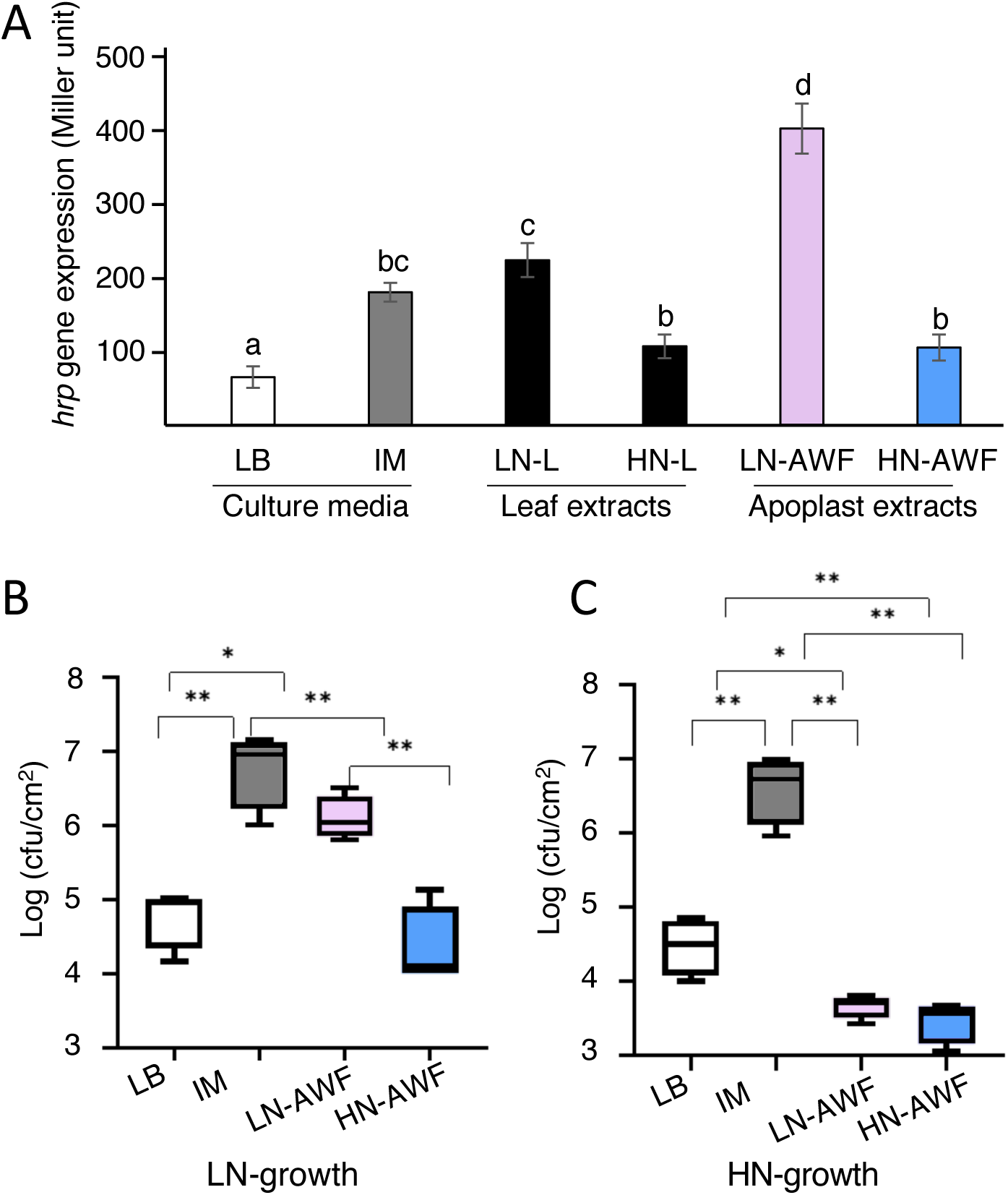
LN-AWF induces high *E. amylovora hrp* expression *in vitro* and increases bacterial fitness s *in planta*. **A**: Apoplast wash fluid (AWF) and whole-leaf extracts were prepared from rosette leaves of plants supplied with 2 (Low N, LN) or 10 (High N, HN) mM NO_3_^-^ *E. amylovora* cells were incubated for 6 h *in vitro* in rich medium (LB), inducing medium (IM), whole leaf extracts (LN-L and HN-L) or AWF (LN-AWF and HN-AWF). *Hrp* gene expression was estimated using the *hrp* promoter-β-Gal reporter fusion. Expression values are given as Miller units as described in Zhang and Bremer (1995). Experiments were performed twice with similar results, the values presented in the graph correspond to the mean of both experiments (n=6). Different letters indicate a significant difference according to the one-way Anova test, post hoc Fisher’s least-significant-difference (LSD) test, P< 0.01. **B, C**: Bacterial titers of wild-type *E. amylovora* in *A. thaliana* leaves supplied with 2 (Low N, LN) or 10 (High N, HN) mM NO_3_^-^. Bacteria were incubated for six hours in LB, inducing medium (IM), LN-AWF or HN-AWF prior to infection. Leaves were sampled 24 hpi and colony forming units (CFU) were measured. B: plants grown in 2 mM NO_3_^-^ : C: plants grown in 10mM NO_3_. Two levels of significance threshold were considered according to the statistical test one-way Anova; p<0.05, one asterisk (*) p<0.05, two asterisks (**) identify adjusted p< 0.01.

To evaluate if different apoplast environments could impact bacterial fitness *in planta*, we incubated *E. amylovora* in two control media, LB and IM, or in AWF-LN and AWF-HN extracts 6 h before leaf infection of plants grown either in 2 mM NO3^-^ (Figure 1 B) or 10 mM NO ^-^ (Figure 1 C). Twenty-four hours post-inoculation (hpi), the bacterial titers *in planta* were measured. Consistently with previous results, *A. thaliana* plants infected with *E. amylovora* preincubated in IM showed a much higher bacterial multiplication than *E. amylovora* preincubated in LB (Figure 1 B and C) (Farjad *et al*. 2021). This strong increase in bacterial titers was independent of N supply. Interestingly, while pre-incubation in HN-AWF did not lead to increased bacterial fitness *in planta* compared to LB, we found that LN-AWF increased bacterial pathogenicity under N limitation (Figure 1B and C).

In conclusion, apoplast extracts isolated from plants grown under N limitation provide an environment that induces *hrp* expression and consequently leads to high multiplication of *E. amylovora* in plants grown under LN. Differences in *hrp* expression between LN-AWF and HN-AWF could be linked to a range of factors, including metabolite content. We therefore analyzed AWF metabolite content to screen for metabolites affected by N supply that could induce or repress *hrp* gene expression.

### N supply alters metabolite content of leaf apoplast

In order to identify the metabolite profiling of apoplast and leaf extracts of plant growth under low and high concentration of nitrogen, we performed GC-MS analysis on whole leaves or leaf apoplast extracts of 5-week-old *A. thaliana* plants grown under low or high N. Our analysis revealed that 146 metabolites were commonly detected in all analyzed samples. Principal Component Analysis (PCA) indicates a clear separation between apoplast and leaf extracts (dimension 1, ∼45%). Yet, a considerable variation was also observed between the apoplast extracts from plants grown with different N supplies (dimension 2, ∼38%; Supp. Figure 1A). To gain more insight into the metabolite composition, we classified the 146 detected metabolites into 7 categories: amino acid, non-polar metabolites, phosphorylated metabolites, fatty acids, organic acids, sugars and others (unclassified metabolites) (not shown). In leaf extracts, the accumulation of these classes of metabolites was similar between LN and HN conditions consistent with the poor PCA separation and our previous observation (Farjad *et al*. 2021). In contrast, all of these classes were less accumulated in LN-AWF in comparison with leaves, indicating an overall lower concentration of metabolite content in LN-AWF extracts compared to leaves (Supp. Figure 1 B). In HN-AWF extracts global sugar, amino acid, organic acid and phosphorylated metabolite levels were similar as in leaf extracts whereas fatty acids and non-polar metabolites accumulated less in HN-AWF than in leaves. Interestingly, the 146 metabolites detected in both apoplast extracts were all found in both LN-AWF and HN-AWF extracts, indicating that none of the detected metabolites accumulated specifically in one N supply condition.

We identified 112 metabolites with significantly different accumulations in LN-AWF and HN-AWF (Figure 2). Several metabolites were strongly accumulated in HN-AWF, such as GABA, citrate and proline for example. Only three metabolites, linolenic acid, levulinate and alpha-tocopherol showed a stronger accumulation in LN-AWF.

**Figure 2:**
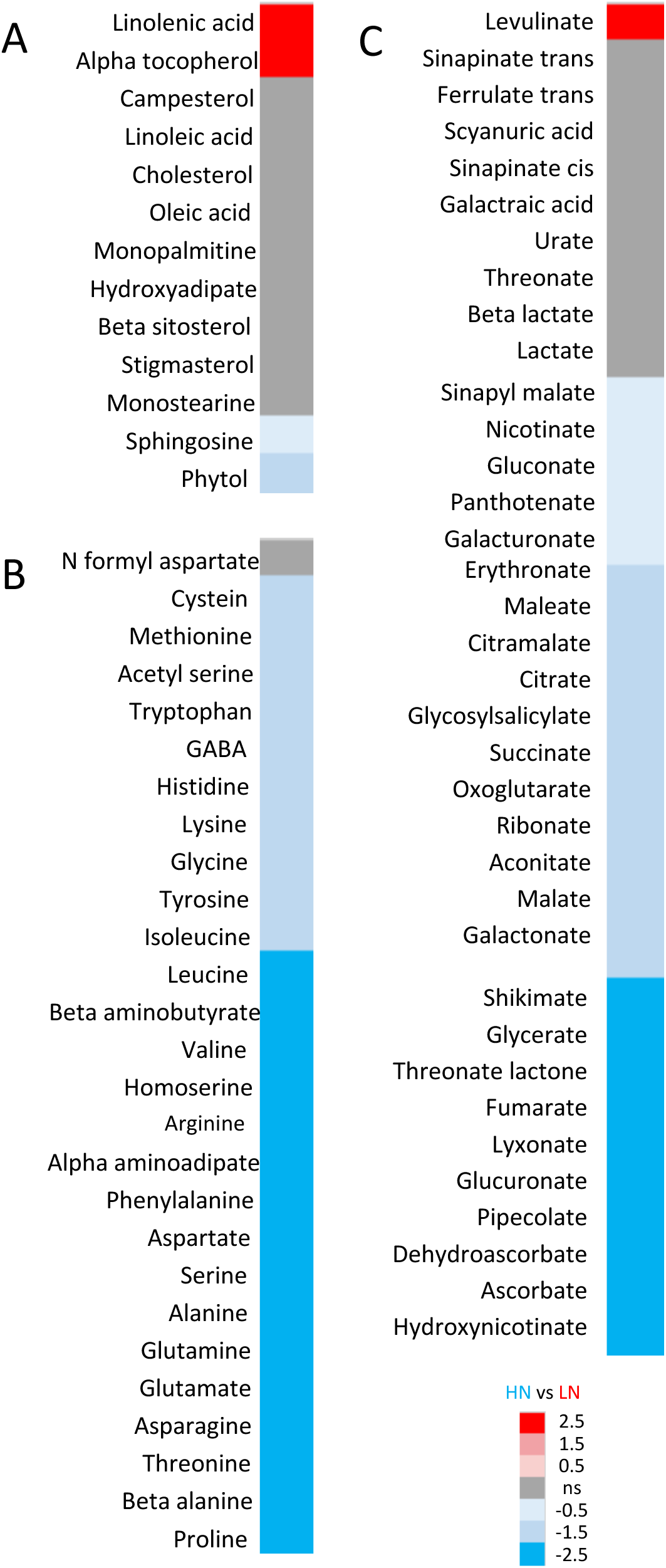
GC-MS analysis of metabolite content in LN-AWF and HN-AWF. Heatmap of metabolites differently accumulated between LN-AWF and HN-AWF. The values correspond to the log2 ratio of metabolites concentration in HN-AWF vs LN-AWF. A: apolar and fatty acids; B: amino acids; C: organic acids. Blue: more accumulated in HN-AWF, Red: more accumulated in LN-AWF, black: non significant. Only selected metabolites are shown.

### Evaluation of the *hrp*-activator and -repressor capacity of selected apoplastic metabolites

Since LN-AWF extracts have *hrp-*inducing properties *in vitro*, we asked if this induction could be caused by specific metabolites accumulated only in LN-AWF. We used the beta-galactosidase activity as a readout to test this hypothesis and incubated *E. amylovora* bacteria in IM medium plus selected metabolites at a standard concentration of 5 mM. Our results show that medium supplemented with the fatty acid linolenic acid and the vitamin alpha-tocopherol triggered a significantly higher *hrp* gene expression than LB (Figure 3A). On the contrary, levulinate, the third most enriched metabolite identified did not show *hrp*-inducing activity as compared to IM medium. We then selected and tested 43 metabolites more enriched in HN-AWF or equally accumulated in both apoplast extracts and found that 29 metabolites had *hrp*-inducing activity *in vitro* (Figure 3B).

**Supplemental Figure 1:**
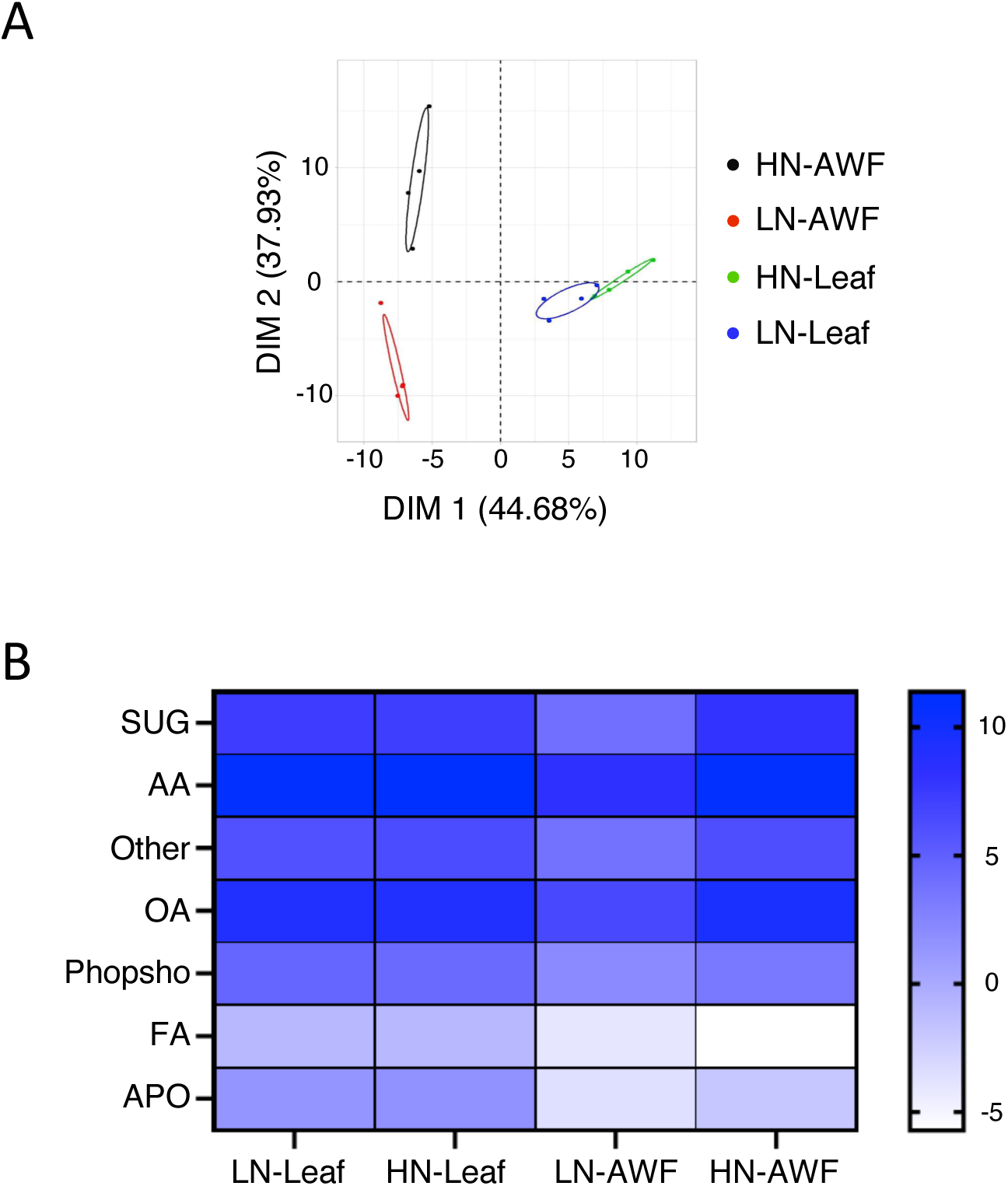
Heat map and PCA of metabolites detected in AWF and leaf extracts. **A**: PCA plot of AWF and leaf metabolite contents in leaf and AWF extracts of plants grown under LN or HN. The dataset corresponds to four replicates per sample. Ellipses represent the Hotteling’s T2 value (95% confidence limit). Samples were separated by PC1 and PC2 which explain 44,68% and 37,93% of the variation between samples, respectively. PCA performed using the Rstudio software. **B**. Heatmap of relative metabolite content in leaf and AWF extracts of plants grown under LN or HN. Plants were grown under low or high nitrate conditions for seven weeks, then an AWF extraction and GC-MS analysis were done. Heat map was based on the mean of four replicates of each metabolite, then the sum of the mean for all metabolites from the same class was done. Presented classes were: sugars (SUG), amino acids (AA), other, organic acids (OA), phosphorylated metabolites (PHOSPHO), fatty acids (FA), and non-polar metabolites (APO). The color of each cell depicts corresponds to the log2 ratio of each metabolite relative to median center metabolite. Ratio=-5 (light blue); ratio =10 (blue). Heatmap generated using Graphpad prism V09.

**Figure 3:**
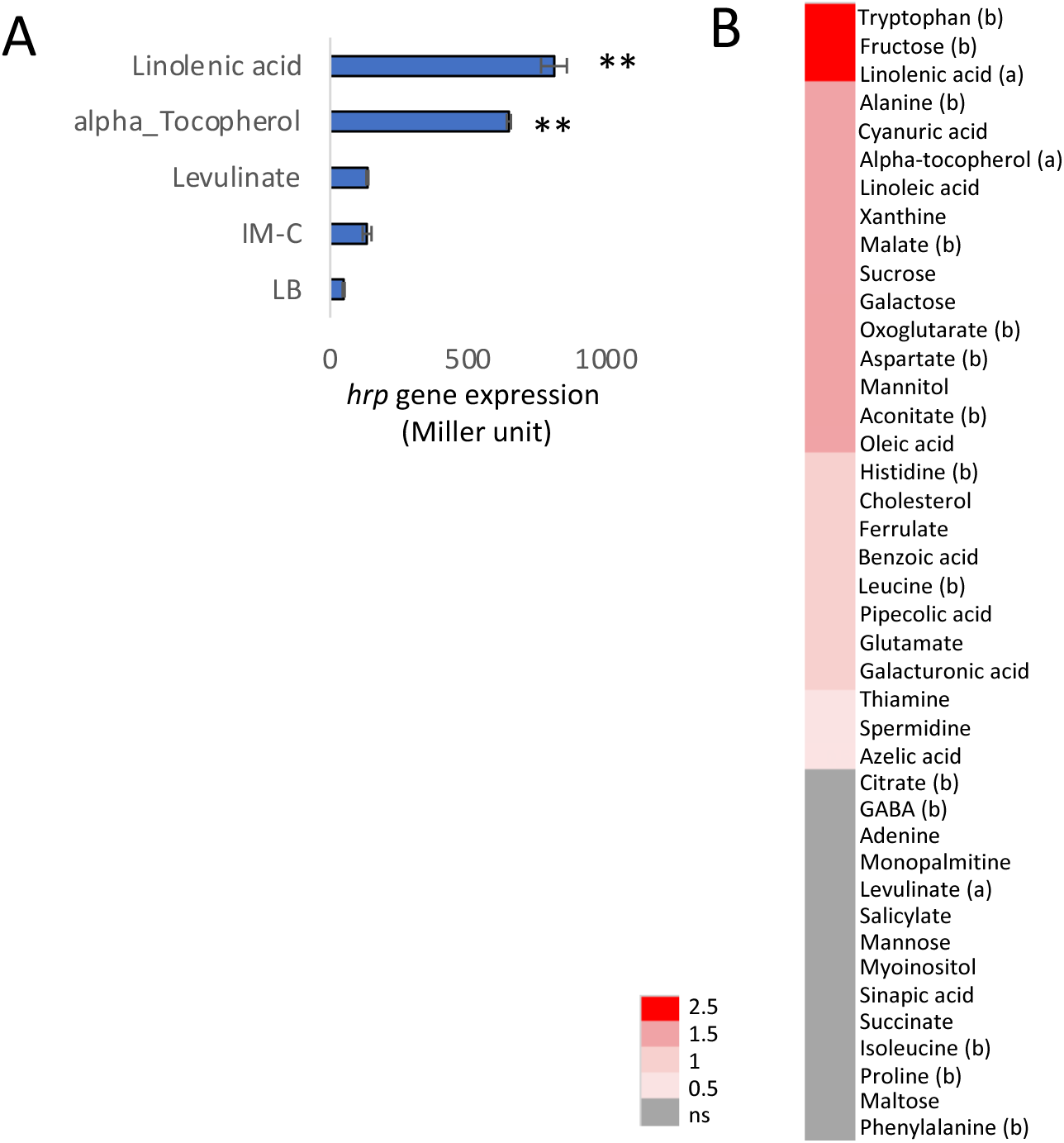
Many metabolites present in the apoplast induce the expression of *hrp* genes. *hrp* gene expression was determined by *LacZ* activity in inducing medium without any carbon source (IM-C), rich medium (LB) and in IM complemented with a single carbon source (as indicated) at 5 mM. The lacZ reporter strain was grown in the corresponding medium at 28°C for 6 h, and beta-GAL activity was measured as described by Jackson and associates (2005). A: we selected the carbon sources among the metabolites found more accumulated in LN-AWF (Figure 1). Stars indicate a significant difference from the LB condition according to the one-way Anova; (P < 0,01). Values shown are the means and standard errors of three replicates. (** p value <0.01). Three independent experiments were performed with similar results. B: Heatmap of induction of *hrp* genes by a larger selection of metabolites, either accumulated in LN-AWF (a), in HN-AWF (b) or not significantly different in the two extracts.

Since several tested metabolites did not trigger *hrp* gene expression, we wondered whether they could display *hrp*-repression activity. We selected two metabolites, citrate and GABA, and tested them in combination with five of the *hrp*-inducers identified in Figure 3, ferrulate, linoleic acid, linolenic acid, cyanuric acid and alpha, to determine whether they can repress *hrp* gene induction. We showed that cotreatment with GABA significantly repressed (p-value < 0.01) the *hrp* gene expression induced by the inducers linolenic acid, cyanuric acid, alpha tocopherol and ferrulate (Figure 4). However, no significant inhibition was observed for cotreatment between GABA and linolenic acid. In addition, we observed that citrate represses the inducer capacity of linolenic acid, cyanuric acid and alpha tocopherol. On the other hand, citrate had no effect on linoleic acid and ferrulate. As a conclusion, both GABA and citrate are *hrp* gene repressors but GABA showed a better *hrp* repressor activity in comparison with citrate.

**Figure 4:**
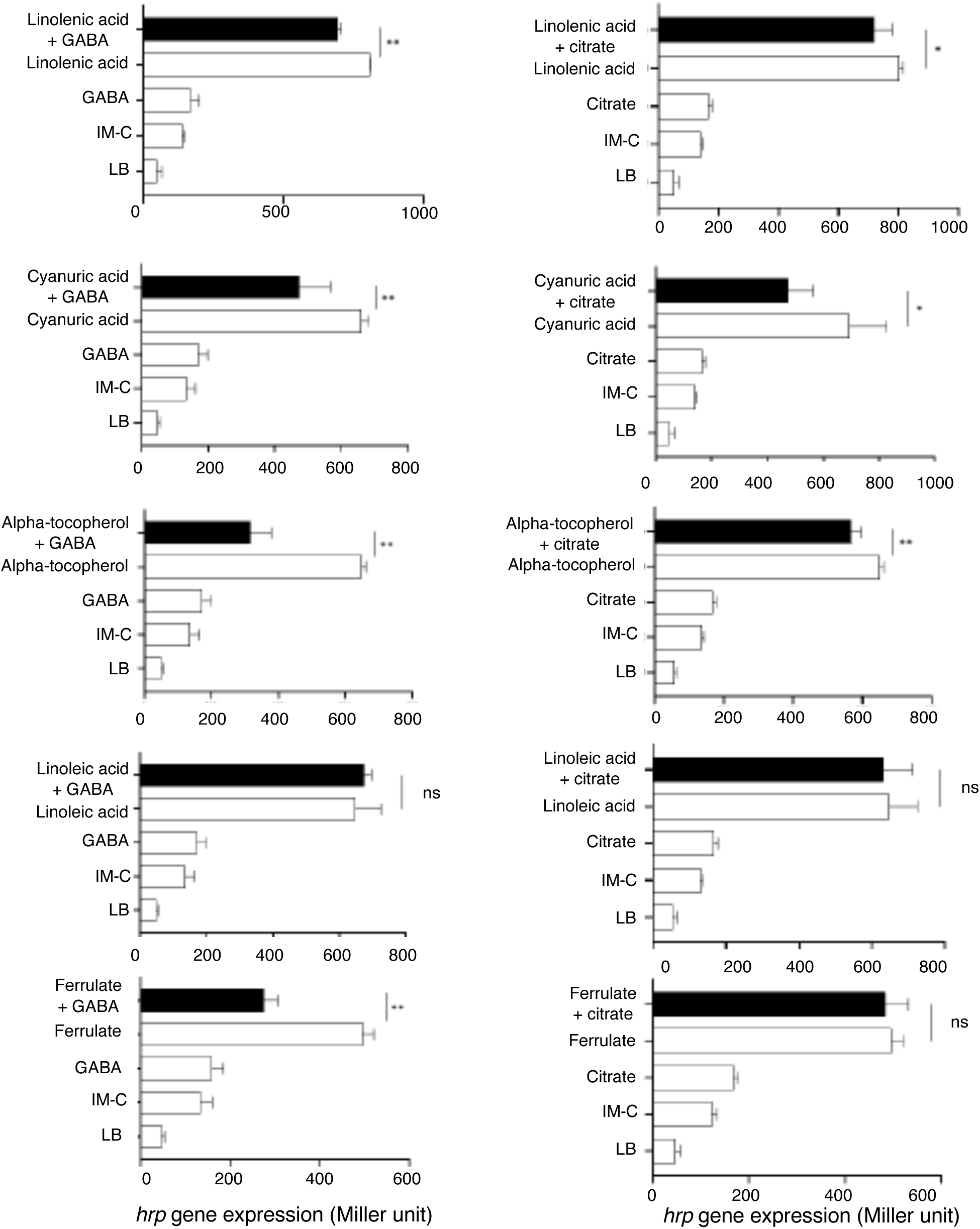
Identification of repressors of *E. amylovora hrp* gene expression. We used the ONPG test to determine *hrp* gene expression. *E. amylovora* was incubated 6H in IM with no carbon source (IM-C) or in IM supplemented with two carbon sources, a potential repressor with a potential inducer. We tested two potential repressors, GABA (left column) and citrate (right column). Each repressor was tested in combination four different *hrp* inducers (black column). All metabolites were at the 5 mM concentration. *hrp* gene expression in the presence of the inducer and the repressor was compared to *hrp* expression in the presence of the inducer alone. Stars indicate significant differences according to one-way Anova test (*: pval<0,05; ** pval<0,01; ns: P > 0.05).

### Bacterial infection impacts apoplast metabolite content earlier in LN conditions

Previously, we identified several apoplastic metabolites able to induce *hrp* genes expression in *E. amylovora* (Figure 3), whereas two had repressive capacity (Figure 4). We wondered how leaf infection by *E. amylovora* affected the apoplastic metabolite content. With this aim, we extracted AWF 6 and 24 h after *E. amylovora* inoculation in plants grown under low or high N supply conditions. As a control, we extracted the AWF from mock-inoculated leaves grown under the same conditions and at the same time-points. GC-MS analysis led to the identification of 91 metabolites. Interestingly, marked differences in the metabolite kinetics were identified for bacterial infection in LN and HN conditions. Indeed, 28 metabolites were modulated as early as 6 hpi, however this was observed almost exclusively in LN-AWF extracts (Figure 5A). Furthermore, in the apoplast extract of infected plants grown in low nitrogen we found GABA, which is able to repress *hrp* gene expression, as well as two inducers (alanine and leucine). At 24 hpi, we found a strong effect of infection by *Ea* on AWF metabolite content in both LN-AWF and HN-AWF (Supp. Figure 2).

**Figure 5:**
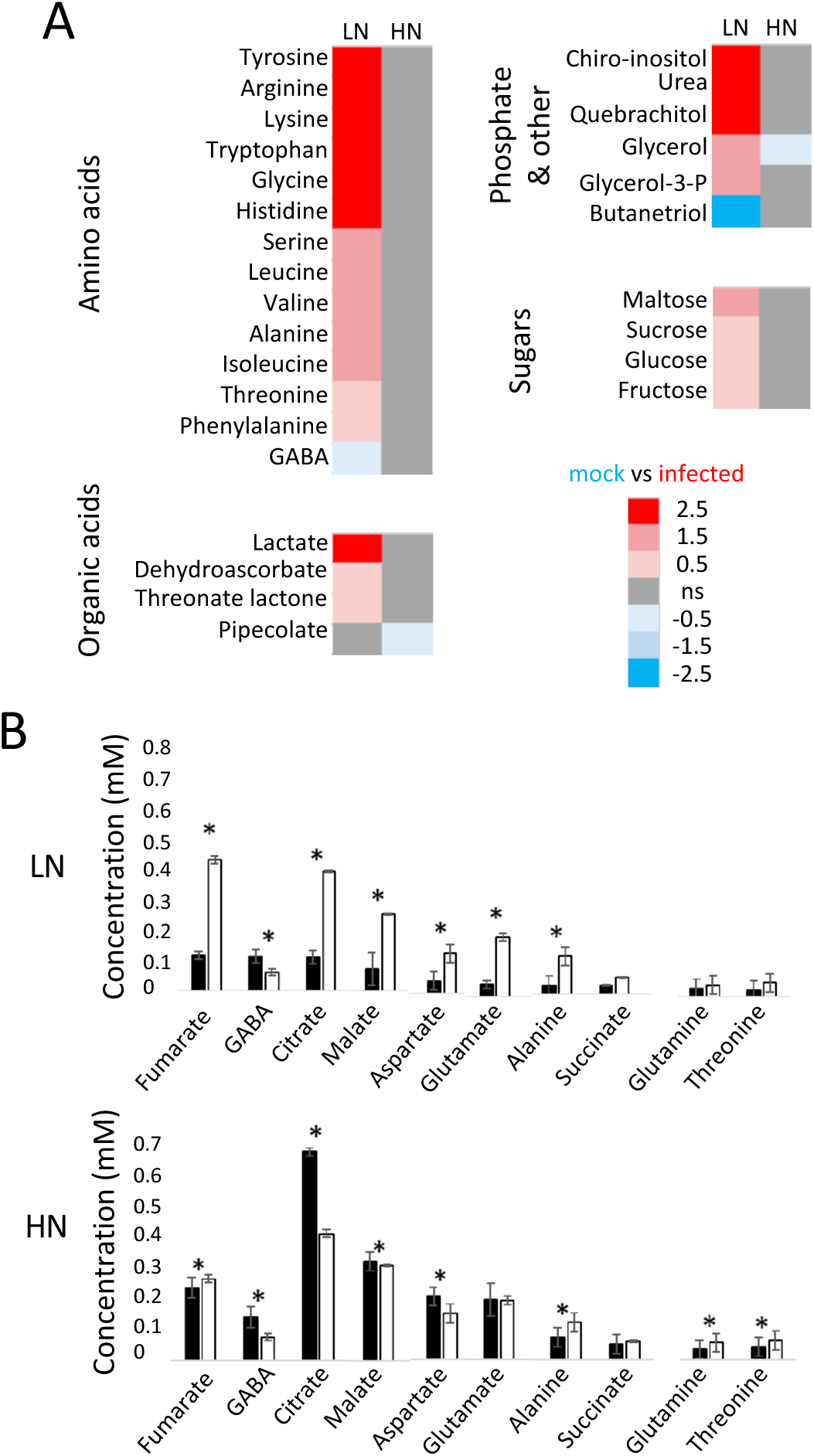
Impact of *E. amylovora* inoculation on metabolite levels in AWF. GC-MS analysis of metabolite content reveals an effect of *E. amylovora* infection on apoplastic metabolome as soon as 6hpi under LN conditions (A) and under both low and high N at 24hpi (B). Metabolite content was analyzed by GC-MS in AWF of *A. thaliana* leaves 6 (A) and 24 (B) hpi following mock inoculation (black bars) or inoculation with *E. amylovora* (*Ea*; white bars). Plants were supplied with 2 (low N, LN) or 10 (high N, HN) mM NO_3_^-^. Error bars represent standard error (n=4). A: heatmap of selected metabolites that accumulate significantly differently between mock and infected tissue at 6hpi. The data corresponds to the log2 ratio of each metabolite content in mock vs infected leaves. Non significant metabolites are shown in grey. B: Quantification of selected metabolites at 24hpi. Stars indicate a significant difference between samples (water and infected samples at 6 or 24 hpi) according to t test (p<0.05).

To determine if bacterial infection specifically modulates metabolites with *hrp*-inducing or - repressing activity, we focused on the most abundant organic acids and amino acids found in the apoplast. We studied their modulation 24 h post bacterial infection in plants grown in low or high N conditions. (Figure 5B). Most of these metabolites were accumulated following infection, indicating that infection had a strong impact on apoplast homeostasis. Interestingly, in LN-AWF metabolites increased following infection, including malate, aspartate and glutamate which are inducers of *hrp* gene expression (Figure 3B). Citrate, which we found to repress *hrp* expression also strongly increased following infection in LN-AWF, while GABA, also a repressor, decreased following infection (Figure 5B). In HN-AWF, most of these metabolites were more accumulated than in LN-AWF before infection as described above, and were only slightly increased or strongly decreased following infection. The most striking effect observed concerns citrate, which is very strongly reduced following infection. Thus, bacterial infection led to a modulation in the homeostasis of the apoplast that is dependent on the abiotic stress endured by the plant.

Altogether, our results show that bacterial inoculation leads to strong modification of AWF content, as early as 6 hpi under LN conditions, including a repression of GABA accumulation. At 24hpi infection led to important modifications in metabolite accumulation including accumulation of several metabolites in LN-AWF.

**Supplemental Figure 2:**
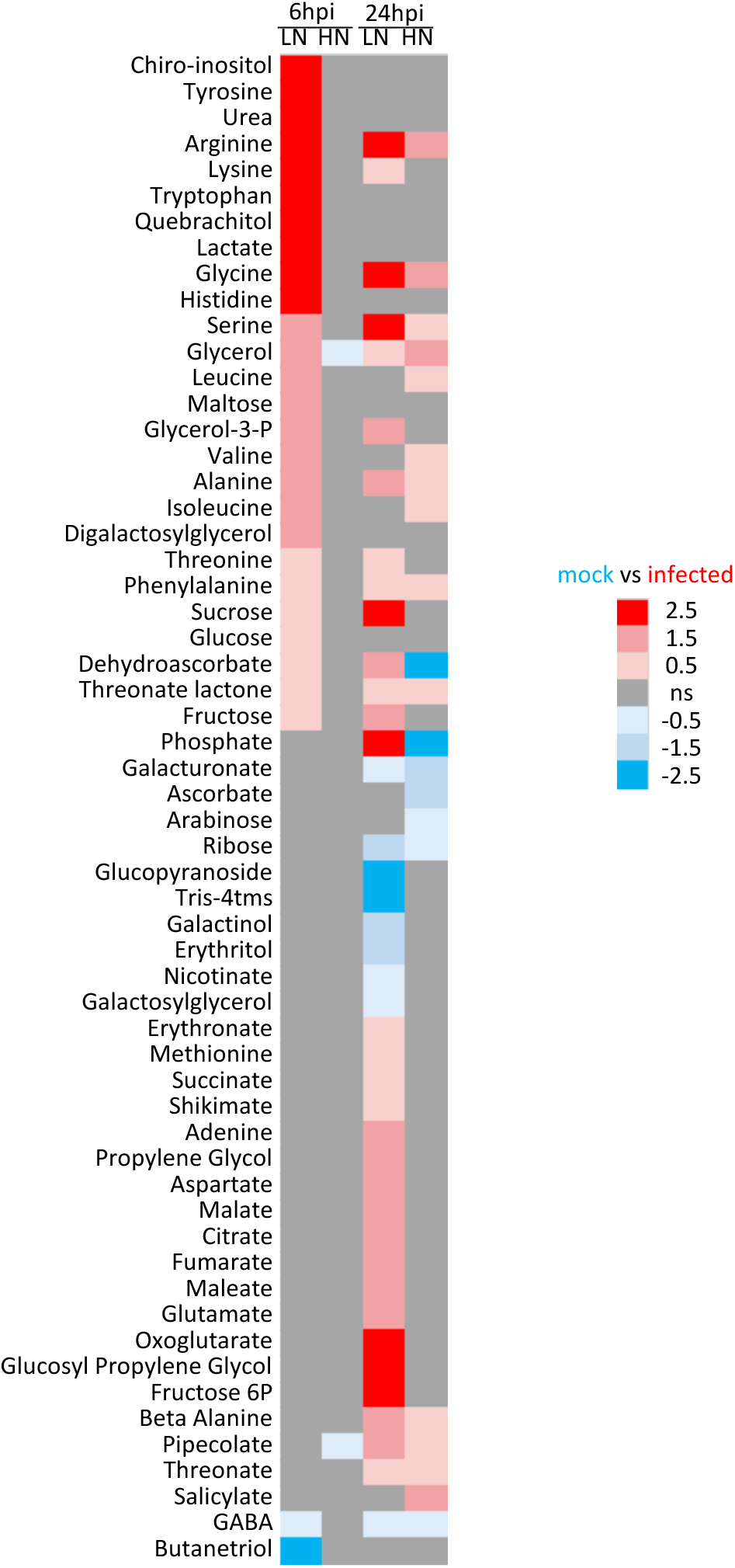
kinetics of impact of *E. amylovora* infection on AWF metabolite content. GC-MS analysis of metabolite content of AWF of leaves following E. amylovora infection. Leaves of plants grown under LN or HN conditions were sampled 6 or 24 hpi following mock or *E. amylovora* inoculation. The results are presented as a heatmap of the log2rat of between mock and infected tissue at 6hpi. Non significant metabolites are shown in grey.

## DISCUSSION

The apoplast corresponds to the plant extracellular space, a crucial compartment in which interactions between plants and microbes take place. The colonization of this compartment is essential for niche establishment by microorganisms. On the contrary, plants counteract niche establishment by modulating the apoplast content as an integral part of plant immunity (Roussin-Léveillée *et al*. 2024). Niche establishment by bacterial pathogens requires manipulation of plant physiological processes to enrich the apoplast in water and nutrients useful to sustain microbe growth. For example, once in contact with apoplastic signals, the timely expression of virulence factors by pathogenic microbes is essential for plant colonization. Although it is established that abiotic stress impacts the outcome of plant-microorganism interactions (Desaint *et al*. 2021; Zarattini *et al*. 2021), the link with niche establishment in the apoplast has not been made.

Numerous studies have described the effect the abiotic stress caused by N limitation on plant physiology, development and plant-pathogen interactions (Fagard et al. 2014; Zarattini et al. 2021). However, to the best of our knowledge, no study has analyzed the impact of abiotic stress on both apoplastic content and its effect on pathogens. Previously, we showed that N limitation in *A. thaliana* increased *in planta* bacterial titers associated with an increase in *hrp* gene expression (Farjad *et al*. 2021). Interestingly, depletion or accumulation of other minerals were shown to increase resistance to pathogens. For example, Fe deficiency leads to an increased resistance to *B. cinerea* (Kieu et al. 2012), while high P leads to increase resistance to several fungal pathogens (Val-Torregrosa et al., 2022). In the case of N limitation, it can lead to increased resistance to *B. cinerea* in *A. thaliana* (Soulié et al. 2021) but reduced resistance in tomato (Lecompte et al. 2010), suggesting that the effect is dependent on the pathosystem. This led us to hypothesize that N limitation could affect apoplastic signals and, consequently, alter the pathogenicity of the bacterial pathogen *E. amylovora*.

To study the impact of N supply on apoplastic content, we isolated apoplast wash fluid (AWF) from *A. thaliana* rosette leaves of plants grown under low N (LN-AWF) or high N (HN-AWF) and performed GC-MS analysis (Figures 2 and 5). We evaluated the capacity of these different extracts to activate *E. amylovora hrp* gene expression *in vitro* (Figure 3). We found that both LN-AWF and HN-AWF displayed *hrp*-inducing activity as was previously reported for the IM minimal medium (Farjad *et al*. 2021; Wei *et al* 1992). Similarly, Xiao and collaborators (1992) showed that *P. syringae hrp* genes are more expressed in leaf tobacco sap extract than in minimal medium. *P. syringae* virulence genes are also induced by tomato apoplast extract although less than in *hrp*-inducing medium (Rico & Preston 2008). The differences observed could be explained by a range of factors, including fluctuations in apoplastic metabolite content, pH variation between plant species, and bacterial nutritional preferences or sensitivity to signals. These results suggest that *hrp* induction is not solely a response to poor medium, as previously suggested, but it is also a pathogenic strategy tightly regulated by several factors including specific metabolites.

In the present study, we found that apoplast fluids extracted from plants grown on low N were stronger inducers of *hrp* genes than those extracted from plants grown in abundant N. We thus compared *in planta* bacterial fitness of *E. amylovora* preincubated in LN-AWF, HN-AWF, and LB and IM known to differentially induce *hrp* gene expression (Figure 1). Our results confirmed that *in planta* multiplication is more important when bacteria are preincubated in IM than in LB medium (Farjad *et al*. 2021). Furthermore, LN-AWF also increased *in planta* bacterial titers as compared to LB-preincubated bacteria (Figure 1). Moreover, bacterial titers were significantly higher when *E. amylovora* was preincubated in LN-AWF than in HN-AWF but only in plants cultivated under LN, indicating that low N is critical in establishing favorable conditions for the interaction. The higher bacterial fitness observed could be due to an enhanced ability of *E. amylovora* to multiply in LN-grown plants and to an increased capacity of LN-AWF-preincubated bacteria to express *hrp* gene. On the contrary, in plants cultivated in high N^-^, preincubation in AWF (LN-AWF or HN-AWF) did not increase *E. amylovora* titers. This confirmed that in high N, bacterial multiplication was not promoted and indicated that it could not be compensated by preincubation in LN-AWF. This latter result suggests that *hrp* induction by the abiotic environment cannot explain all of the results observed in plants grown in LN. This suggests either that other mechanisms could be triggered by IM. Alternatively, LN-AWF beyond being an inducing environment for *hrp* expression, could enhance *in planta* bacterial multiplication by other means, also dependent on the environment.

Given the higher capacity of LN-AWF to induce *E. amylovora hrp* induction and the low capacity of HN-AWF to do so (Figure 1), we hypothesized that either LN-AWF contained more metabolites with *hrp*-inducing activity than HN-AWF or that HN-AWF contained more metabolites with *hrp*-repressing activity. To test this hypothesis we performed a GC-MS analysis of AWF extracts obtained either from rosette leaves of plants grown under low or high N. Consistently with previous reports, we observed that the metabolite content of AWF extracts was distinct from the leaf extracts (Freire, Morais & Daloso 2024). Indeed, although the same metabolites were found in leaf and AWF extracts, their relative abundance differed in the leaf and in the AWF, especially for organic acids, amino acids and fatty acids (Supp. Figure 1). Furthermore, we compared the relative amounts of metabolites in LN-AWF and HN-AWF and found important differences, with 109 metabolites significantly more accumulated in HN-AWF whereas three were more abundant in LN-AWF (Figure 2). We then evaluated the *hrp*-inducing capacity of 43 selected metabolites and found that 28 of them acted as *hrp*-inducers (Figure 3). Among the metabolites able to induce *E. amylovora hrp* genes we found sugars, for example mannitol, fructose and sucrose, which are carbon sources abundant in the apoplast, but also organic acids and amino acids, some of which are known to be used by *E. amylovora* as C and N sources. Likewise, we tested the capacity to repress *hrp* gene expression *in vitro* of two metabolites, citrate and GABA, found more accumulated in HN-AWF (Figures 1 and 3). Interestingly, our results showed that GABA and citrate are both repressors of *E. amylovora hrp* genes *in vitro*. This observation is consistent with a previous study in which both metabolites were shown to inhibit *P. syringae hrp* genes (Park *et al*. 2010).

Bacterial *hrp* genes, which encode the type III secretion system (T3SS), a delivery apparatus for type three effectors used by many pathogenic bacteria to establish disease (Asif, Xie & Zhao 2024). What causes the induction of *hrp* genes *in planta* is nowadays still not completely understood. Several hypotheses have been proposed including an alteration of physical barriers such as the cell wall (Xiao *et al*. 2004), pH variation or the presence of inducing metabolites (Khokhani *et al*. 2013; Anderson *et al*. 2014). Our results clearly show that several apoplastic metabolites, accumulated following abiotic stress, can modulate *hrp* gene expression in *E. amylovora* (Figure 3). Although we observed that LN-AWF has higher *hrp*-inducing capacities than HN-AWF, we could not correlate this with specific metabolites. However, we found that two metabolites, citrate and GABA, had *hrp*-repressing activity which could explain the low *hrp*-inducing activity of HN-AWF.

Previously, several metabolites were shown to influence *hrp* gene expression in pathogenic bacteria. For example, oleanolic acid (Wu, Ding, Zhang, Liu & Yang 2015), coumaric acid, caffeic acid cinnamic acid, and ferrulate (Zhang *et al*. 2017), pyroglutamic acid, citrate and shikimate (Anderson *et al*. 2014), o-coumaric acid, t-cinnamic acid and p-coumaric acid (Khokhani *et al*. 2013) were shown previously to induce *hrp* genes in *P. syringae*, *R. solonacearum*, or *E. amylovora*, respectively. On the contrary, several apoplast-abundant amino acids, and tricarboxylic acid (TCA) intermediates were shown to repress *hrp* gene expression. For example, thiazolidinone (Wu, Ding, Zhang, Liu & Yang 2015) and GABA repress *P. syringae hrp* gene expression in tomato (Park *et al*. 2010). Also, citrate and succinate block *P. syringae hrp* gene expression in tobacco and bean (Rahme *et al*. 1992). Recently, 4-methylsulfinylbutyl isothiocyanate, a natural product derived from an aliphatic glucosinolate called glucoraphanin, was identified as a repressor of *P. syringae hrp* genes (Wang et al. 2020). Interestingly this metabolite is also involved in the control of plant immune responses such as callose deposition and programmed cell death (Piasecka, Jedrzejczak-Rey & Bednarek 2015). In the present study, we found that 28 metabolites show *hrp*-inducing activity *in vitro* (Figure 3), several of which, such as linolenic, linoleic and oleic acids, had not been previously identified as *hrp* inducers.

To deepen our investigations, we analyzed the effect of *E. amylovora* infection on apoplast content to see how the concentration of *hrp* inducers and repressors fluctuates in response to infection in low and high N. To evaluate the effect of infection with *E. amylovora* on the AWF content, we performed GC-MS analysis and detected 91 metabolites. We observed that the AWF of plants grown under low N and infected with *E. amylovora* had an altered metabolite content as early as 6 hpi, mostly due to an increase in the concentration of several metabolites (Figure 5 and Supp. Figure 2). For example, we observed an increase in several sugars, such as sucrose, glucose and fructose, in LN-AWF 6 h following infection, a sign of successful infection. Moreover, we found that a large number of metabolites increased 24h following inoculation while only a few metabolites were depleted 24h following infection. Interestingly, the metabolomic profile of LN-AWF and HN-AWF 24h following infection were very different, supporting the importance of the environment on the infection process at the apoplastic level. Furthermore, at this later timepoint we observed an increase in salicylate, a key regulator of plant defenses. Concerning the differences in the kinetics of modification of AWF by infection between LN and HN conditions, they could be explained by the high bacterial titers observed in plants grown in low N. Previous studies have shown an important effect of infection on plant apoplast content. For example, the infection with the fungus *C. fulvum* in tomato leaf increases the concentration of mannitol, GABA, valine, tyrosine, proline, leucine, isoleucine, lysine, and arginine in the apoplast (Solomon and Oliver 2002). Also, *B. cinerea* and *P. syringae* infection led to a decrease in the level of apoplastic sugar (Berger, Sinha & Roitsch 2007).

Different mechanisms could alter metabolite levels in the apoplast of plants under biotic stress. One of them is apoplast alkalization caused by infection, which could modify the transport of metabolites through the plasma membrane and lead to nutrient accumulation in the apoplast. Moreover, the T3SS of pathogenic microorganisms can create pores in the plasma membrane through which effectors penetrate into the cytoplasm and cause an efflux of cations from the cytosol to the apoplast (Engelhardt *et al*. 2009). In turn, the decrease of membrane polarization then leads to the accumulation of cytosolic metabolites in the apoplast content as the export of citrate with K^+^ efflux to maintain the electric charges. Also, infection leads to an increase of sucrose efflux and the membrane sugar transport protein (SWEETs) in *A. thaliana* due to the increase of pH and K^+^ efflux (Chen 2014).

In summary, in our study we screened primary metabolites present in the LN-AWF and HN-AWF and identified that citrate and GABA have an inhibitory potency on *hrp* genes and that 27 metabolites have *hrp*-inducing capacities. Previous studies have demonstrated the importance of the environment for the expression of virulence genes in plant pathogenic bacteria. Since the apoplast is one of the primary interfaces of plant-pathogen interactions, its manipulation could be an alternative way to increase plant resistance. For example, increased GABA and citrate content in the apoplast could lead to reduced susceptibility to bacterial pathogens. Interestingly, GABA often accumulates in leaves in response to biotic stress, such as bacterial pathogens (Li *et al*. 2021; Farjad *et al*. 2021). Although relatively few is known about GABA secretion in the apoplast, a transporter has been identified in wheat (Ramesh *et al*. 2018). Further investigation concerning the homeostasis of GABA in the apoplast in response to multistress would thus be very interesting.

## MATERIAL AND METHODS

### Plant material and growth conditions

Experiments were performed on *A. thaliana* plants accession Col-0. Plants were grown in a culture chamber with 8 h light at 21°C and 16 h night at 18°C with 150 µE/m2/s irradiation and 65% humidity. Plants were watered three times a week with a nutrient solution containing 2 mM NO_3_ ^-^ (low N) or 10 mM NO_3_ ^-^ (high N) as described previously (Soulié et al 2020; Farjad *et al*. 2021).

### Bacterial strains and culture media

For inoculation experiments, we used the previously described wild-type strain of *E. amylovora* CFBP1430 (Farjad *et al*. 2021). For *hrp* gene expression assays, we used the previously described *E. amylovora* CFBP6023 strain carrying the *lacZ* reporter gene downstream of the *hrp* promoter (Farjad *et al*. 2021).

For all experiments, bacteria were initially grown on solid Luria Broth (LB) w/ agar (15 g/l) at 28°C for two days and overnight in liquid LB at 28°C, shaken at 200 rpm. To induce *E. amylovora hrp* gene expression we used the previously described inducing medium (IM): M9 salts (pH 7.4), 0.2% nicotinic acid, 2 % galactose (Wei *et al*. 1992; Farjad *et al*. 2021) or another carbon source as indicated (Figures 3 and 4).

### Assay of *hrp* gene expression using fJ -galactosidase activity

To assay *hrp* gene expression in the *E. amylovora* reporter strain carrying the lacZ reporter gene downstream of the *hrp* promoter, the galactosidase assay was carried out based on previous studies with slight modifications (Zhang & Bremer 1995). After overnight incubation, the OD600 of the bacterial cultures were diluted to 0.8/0.9. Then, 1 mL of culture was washed twice with sterile water followed by centrifugation, to remove all traces of culture medium. The resulting pellet is resuspended in 600 µl of sterile distilled water (dH2O). To induce *hrp* gene expression, 200 µl of the bacterial suspension is placed in 200 µl of IM, LB, LN-AWF or HN-AWF. Bacteria are then incubated for 6h, then washed twice with water, and centrifuged. The recovered pellet is resuspended in 100 µl dH2O. Then, 20 µl of bacteria were taken and 80 µl of permeabilization solution (100 mM Na2HPO4, 20 mM KCl, 2 mM MgSO4, 0.8 mg/ml CTAB, 0.4 mg/ml sodium deoxycholate, 0.27 mM B-mercaptoethanol) is added, followed by 600 µl of substrate solution (60 mM Na2HPO4, 40 mM NaH2PO4, 1 mg/ml ONPG, 0.135mM B-mercaptoethanol). The samples are incubated at 30°C until a yellow color appears, then the reaction is stopped by adding 700 µl of stop solution (1 M Na2CO3). Then 200 µl of supernatant is transferred to a transparent 96-well plate and absorbance at 420 nm is measured using a Tecan Spark 10M plate reader (Austria).

### Leaf inoculation

For bacterial inoculum preparation, an aliquot of the glycerol stock culture was streaked onto agar medium and grown for 48 h at 28°C. Bacteria grown on Petri dishes were used to inoculate a liquid culture under agitation at 28°C. After 8 h of growth, 100 µl of the culture was spread on agar medium without antibiotics and allowed to grow overnight at 28°C. A bacterial suspension of *E. amylovora* with an OD of 0.1 (10^7^ CFU/ml) in water was inoculated into 7-week-old plants using a syringe.

### Extraction of apoplastic fluid

Apoplast fluid was extracted using the infiltration-centrifugation method with slight modifications (O’Leary *et al*. 2014). The resulting extracts are referred to as apoplast wash fluid (AWF) extracts. For each point 38 to 40 *A. thaliana* leaves from 6 individual 7-week-old plant are used. The detached leaves are placed in 200 ml of milliQ water and submitted to two pressure cycles of -25 Pa vacuum using a vacuum pump. The infiltrated leaves are then gently dried with paper, rolled in Parafilm and inserted into a 20mL syringe. The syringe is then placed inside a 50 mL conical tube. Centrifugation of the syringe placed in a conical tube at 1000g for 20 min at 4°C collects the AWF samples. The resulting AWF is centrifuged again for 10 min at 1000 g/10 min at 4°C to remove cellular debris. The supernatant is collected in a new 1.5 ml tube.

### Measurement of AWF cytoplasmic contamination

To estimate the presence of intracellular contaminants in the AWF, we compared Malate dehydrogenase (MDH) activity in AWF and whole leaf extracts as previously described (O’Leary *et al*. 2014). MDH activity was determined spectrophotometrically (340 nm) at 25°C in a medium containing 0.2 ml of MOPS buffer pH 7.5 (0.1 M), 0.05 ml of NADH (0.5 mM), 0.01 ml of whole leaf extract or AWF, and 0.54 ml dH2O in a total volume of 1 ml. The reaction was started by adding 0.2 ml of 2 mM oxaloacetic acid. The activity is measured as the OD340 per minute. In parallel, we also measured chlorophyll contamination of AWF extracts. After centrifugation, AWF was removed and a pellet of chlorophyll remained in the tube which was dissolved in 750 ml of ethanol (95%). Then OD at 664nm was measured and compared to the leaf extract chlorophyll content.

### Estimation of AWF dilution

To estimate the dilution of metabolites in AWF we proceeded as described in Baker *et al*. (2012). Briefly, two sets of leaves are used for each condition, the first set is infiltrated with water, the second set is infiltrated with a solution of indigo carmine (indigo 5,5’-disulfonic acid 50 mM pH 6.2). A conventional AWF extraction is then performed with both solutions and the OD is measured at 610 nm. Next, the dilution factor is calculated using the following equation: dilution factor = OD AWFIC / (OD AWFIC - OD AWFE), where IC= indigo carmine; E = water (O’Leary *et al*. 2016).

### Metabolome analysis

Metabolome of AWFs (see above) was performed by GC-MS as previously described (Farjad *et al*. 2021). As a control, metabolome of whole leaves was analyzed. For each condition, 7-8 whole leaves were collected in liquid nitrogen and then ground to serve as a control. Between 40 and 50 mg of frozen powder was processed through an Agilent 7890A gas chromatograph (GC) coupled to an Agilent 5975C mass spectrometer (MS). For quantification, standards were injected at the beginning and end of the analysis. Data were analyzed with AMDIS12 and QuantLynx software (Waters). Metabolome analysis was done using the concentration of each metabolite to total metabolites [µg/ (ml)]. For statistical analysis we used Graphpad prism 9 and R studio software.

## ACKNOWLEDGEMENTS

This work has benefited from the support of IJPB’s Plant Observatory platforms PO-Plants and P0-Chem. The IJPB benefits from the support of Saclay Plant Sciences-SPS (ANR-17-EUR-0007). MZ was supported by an FNRS postdoctoral grant. We thank R. Berthomé, J. Colcombet, M. Delarue, D. Expert, S. Jasinski and HN Truong for fruitful discussions.

